# Label-Free Tracking of Subcortical White Matter Degradation *In Vivo* Using Third Harmonic Generation Microscopy in a Mouse Model of Multiple Sclerosis

**DOI:** 10.1101/2025.07.31.667796

**Authors:** Nicole E. Chernavsky, Nuri Hong, Lianne J. Trigiani, Nozomi Nishimura, Chris B. Schaffer

## Abstract

Early detection and characterization of myelin degradation in the white matter (WM) is important for understanding neurodegenerative diseases such as multiple sclerosis (MS) and dementia. Here, we demonstrate a label-free *in vivo* imaging technique using third harmonic generation (THG) microscopy at 1320-nm excitation, which enables visualization of subcellular myelin structural changes deep in the mouse brain. By applying longitudinal THG imaging of the same axons in the cuprizone mouse model of MS, we captured the formation, evolution, and regression of myelin blisters in the corpus callosum through intact cortex - a depth inaccessible to other optical methods. We also confirmed the utility of conducting THG imaging in parallel with three-photon excited fluorescence imaging, which allowed us to determine that myelin blistering events were not correlated with the location of microglia cell bodies. Further, we used post-mortem immunohistochemistry to establish the ability to identify and measure the intranodal distance at nodes of Ranvier using *in vivo* THG imaging, and we found increased intranodal distances after cuprizone administration. We also developed a novel quantitative metric based on spatial concentration of the brightest THG signal that occurs before more overt changes like myelin blistering. Overall, THG microscopy offers a powerful method for detailed tracking of subcortical myelin dynamics, providing new opportunities to investigate disease mechanisms and potential therapeutic interventions in MS and other neurodegenerative diseases.

## 1 Introduction

Dementia, multiple sclerosis (MS), and other neurodegenerative diseases are devastating for patients and their families. Unfortunately, once these diseases are diagnosed, treatment options are limited, so the need to identify and understand early-stage biological processes and biomarkers is critical. To this end, white matter (WM) changes have been identified both as a biomarker of degeneration as well as a potential contributing factor to brain dysfunction in many neurodegenerative diseases (Festa et al., 2024). Recent studies of MS have sought to understand the early degenerative changes that occur in focal WM lesions, such as myelin blistering (Luchicchi et al., 2021; Hart et al., 2021), and the detailed cellular dynamics that contribute to these lesions remain an active area of investigation (Gay et al., 1997; Barnett et al., 2004; Kutzelnigg et al., 2005; Marik et al., 2007; Lassmann, 2018; Luchicchi et al., 2024).

To better understand this myelin degradation in the subcortical WM of MS and other diseases, the ability to observe myelin changes on a microscopic scale in live animal models is essential. Specifically, following myelin structure over time in mouse models could help uncover mechanisms of myelin degradation and potential treatments. To date, several techniques have been explored to visualize myelin *in vivo*. T2-weighted magnetic resonance imaging (MRI) and diffusion tensor imaging are useful for visualizing the entire brain, mouse or human, but they are limited to millimeter-level spatial resolution that cannot capture detailed structural changes (Fox et al., 2011; Enzinger et al., 2015). Coherent anti-Stokes Raman scattering (CARS) microscopy, on the other hand, can measure myelin organizational changes *in vivo* on the sub-micrometer scale by using two laser beams to spectroscopically visualize the vibrations of C-H_2_ bonds, which are abundant in lipid-rich materials like myelin (Cheng, 2007; Fu et al., 2008; Yu et al., 2014). Stimulated Raman scattering (SRS) can similarly highlight lipid-rich features based on molecular vibrations, but without the nonresonant background of CARS (Yu et al., 2014). Polarization-resolved CARS can also obtain both microscopic and molecular structure of the myelin sheath *in vivo* by measuring the alignment of the C-H_2_ molecular dipoles (Turcotte et al., 2016). Yet, as useful as CARS and SRS have become for imaging myelin *in vivo*, these techniques are limited by imaging depth and would therefore not translate well for imaging subcortical WM in a mouse model. They are also somewhat complicated, requiring two spectrally tuned, temporally synchronized laser pulse trains overlapped in the microscope. Another technique - spectral confocal reflectance microscopy (SCoRe) - can be used to identify sub-micrometer features in myelin (Schain et al., 2014) up to 400 µm deep into the mouse cortex. Unfortunately, this is also still less than half the depth needed to study the subcortical WM in mice at 800 µm and below.

Meanwhile, third harmonic generation (THG) microscopy is a technique that has shown promise in visualizing myelin *ex vivo* and *in vivo* at the necessary depth and resolution to track structural changes. THG is a nonlinear optical process where three photons are coherently converted to a single photon at one-third of the initial wavelength, propagating in the forward direction (Boyd, 2008). In the case of 1320-nm excitation, THG is thus produced at 440 nm. In a uniform material, the THG signal is not visible in the far-field because the THG generated on either side of the focus is out of phase due to the Gouy phase shift experienced by a Gaussian beam passing through a focus. When an interface disrupts the symmetry around the focus, however, imperfect cancellation of THG signal generated on either side of the focus leads to an increase in the THG detected in the far field (Barad et al., 1997; Squier et al., 1998). This effectively makes THG an interface-sensitive imaging technique, wherein changes in the refractive index or the nonlinear susceptibility produce THG image contrast at the interface (Tsang, 1995; Clay et al., 2006). Our group has used this technique of label-free imaging to identify atherosclerotic plaques (Small et al., 2017), granules in Paneth cells in the gut (Choi et al., 2018), flowing red blood cells in mouse cortex (Ahn et al., 2020), and myelinated axons in the spinal cord (Farrar et al., 2011) - all of which have bold optical interfaces. In particular, the multiple lipid layers of myelin sheath result in a relatively thick interface with a very strong THG signal that increases exponentially with the thickness of the myelin sheath (Farrar et al., 2011).

Although prior work showed bright THG signals from myelinated axons in the WM of the mouse spinal cord (Farrar et al., 2011), the necessary depth for subcortical WM imaging could not be reached, due to the use of 1034-nm excitation. As femtosecond laser systems have been developed at longer excitation wavelengths, primarily to facilitate three-photon excited fluorescence (3PEF) microscopy (Horton et al., 2013; Prevedel et al., 2025), this has enabled THG microscopy to also be performed at greater depths. In this paper, we optimized our label-free THG microscopy technique at 1320-nm to acquire three-dimensional high-resolution structural images of subcortical WM, through the intact cortex, in live mice. We then applied this technique to the cuprizone mouse model of MS and observed dynamic changes in myelin structure over time, thereby paving the way for the exploration of myelin structural dynamics in a brain region that is relevant for many neurodegenerative diseases.

## 2 Materials and methods

### 2.1 Animals

Four transgenic mouse strains as well as C57BL/6 wild-type (WT) control mice were used in this study. (1) Mice with a targeted replacement of murine ApoE with human ApoE3 (B6.Cg- *Apoe^em2(APOE*)Adiuj^*/J – Jackson Labs; #029018) were bred from heterozygous to homozygous on site and were used, along with WT mice, to represent a normal WM architecture during development of the imaging technique. (2) CX_3_CR-1^GFP^ knock-in/knock-out mice (B6.129P2(Cg)- *Cx3cr1^tm1Litt^*/J – Jackson Labs; #005582) were bred on site and used to track microglia activation in the subcortical WM. (3) One male PLP-EGFP mouse (B6;CBA-Tg(Plp1-EGFP)10Wmac/J – Jackson Labs; #033357) was used to observe oligodendrocyte morphology in the subcortical WM. (4) One Thy-1 YFP-16 mouse (B6.Cg-Tg(Thy1-YFP)16Jrs/J - Jackson Labs; #003709) was also used to visualize axon-myelin dynamics *in vivo*. Male and female C57BL/6 WT and other transgenic mice were aged 6-12 months in experiments. This study was carried out in accordance with the recommendations of Guide for the Care and Use of Laboratory Animals by the National Institutes of Health. The protocol was approved by the Institutional Animal Care and Use Committee of Cornell University (protocol #: 2015-0029). Supplementary Table 1 shows which mice were used for each study in this manuscript.

### 2.2 Chronic cranial window preparation

To achieve optical access to the subcortical WM, the skin and skull were surgically removed and replaced with an 8-mm diameter glass coverslip (0.17 mm thickness, Electron Microscopy Sciences #72296-08). For the procedure, the mice were anesthetized with isoflurane (1.5-2% in oxygen), then given glycopyrrolate (0.5 mg/100 g animal weight; intramuscular; #00143968125, Hikma) to mitigate fluid buildup in the lungs and 5% weight/volume (w/v) glucose in saline (200 μL; subcutaneously; D-glucose, Sigma-Aldrich) to provide hydration and energy during anesthesia. Body temperature was maintained at 37°C via a feedback-controlled heating pad (50-7053P; Harvard Apparatus). A bilateral craniotomy over the parietal cortex, between the lambda and bregma sutures, was performed using a 0.7-mm dental drill, as described in Holtmaat et al. (2009). Sterile surgical foam (Surgifoam #1972, Ethicon) was used to alleviate bleeding as needed. Once the skull was removed and any bleeding had subsided, the exposed brain was covered with a drop of sterile saline and the 8-mm glass coverslip was placed over the opening. The coverslip was sealed in place with cyanoacrylate glue (Loctite 495; Henkel) and dental cement (Co-Oral-Ite Dental Mfg Co.). For three days following the procedure, mice recovered with a heating pad under half of their cage and were given daily dexamethasone (0.025 mg/100 g; intramuscular; 07-808-8194, Phoenix Pharm Inc.) and ketoprofen (0.5 mg/100 g; subcutaneous; Zoetis Inc.) subcutaneously to reduce pain and manage inflammation.

### 2.3 Third harmonic generation microscopy setup

THG and 3PEF were excited using 1320-nm laser light generated by an optical parametric amplifier (OPA) (Opera-F, Coherent) that was seeded with a diode-pumped femtosecond laser (40 μJ/pulse at 1 MHz; Monaco, Coherent). The resulting pulses have a duration of ∼50 fs and ∼1.3-μJ energy at 1-MHz repetition rate. The laser power was controlled by rotating a polarizing beam splitter (PBS254, 1200-1600 nm, Thorlabs) and dispersion compensation was achieved with a SF11 prism pair, optimized for maximum 3PEF signal at the sample. Maximum power at the objective during *in vivo* imaging was ∼200 mW to avoid thermal damage to the sample (Podgorski, Ranganathan, 2016). To generate an image, the laser was raster scanned with galvanometric scanners (bi-directional; Saturn 1 System 4-mm, ScannerMax) at a line rate of ∼0.5 kHz and a 3.2-μs pixel dwell time, resulting in 2-3 laser pulses per image pixel. From the scan mirrors, the laser passed through an achromatic scan lens (80-mm focal length; AC508-080-C, Thorlabs) and an achromatic tube lens (300-mm focal length; AC508-300-C, Thorlabs) before entering the back aperture of a 25x objective (XLPlan N 1.05 NA, Olympus).

Excitation light and emitted THG/3PEF light were separated by a long-pass dichroic mirror (720-nm cutoff, Semrock), and the THG/3PEF light were further separated by 473-nm and 593-nm dichroic mirrors into three channels and focused via collection lenses onto the 5-mm active area of the GaAsP photomultiplier tubes (H7422P-40, Hamamatsu) used as detectors (Tsai et al., 2015). Signals from each photomultiplier tube were passed through custom pre-amplifiers (gain of 20 and 10-MHz bandwidth) and 500 kHz low-pass filters (EF506, Thorlabs). Using ScanImage software (version 2022.0.0, Vidrio Technologies) to control laser power, scanning, and stage movement (Pologruto, Sabatini, Svoboda, 2003), three channels of data with 512-by-512 pixels per frame were acquired with this setup. The emission channels were additionally filtered as follows: THG with 447/60 nm band-pass filter (BPF), green fluorescence with 540/80 BPF, and red fluorescence with 630/92 BPF (center wavelength/bandwidth for BPF).

### 2.4 Cuprizone diet

Following the protocol in Zhan et al (2020), mice were fed a diet of 0.2% cuprizone (bis(cyclohexanone)oxaldihydrazone) (#C9012; Sigma Aldrich) for up to 8 weeks. The diet was prepared fresh every other day by grinding the standard rodent chow (Envigo 7912) to a fine powder using a blender and then a flour mill. The appropriate amount of cuprizone was weighed out and mixed into the powdered food. During feeding, the mice were also weighed to monitor their health and consumption of the cuprizone diet. After 8 weeks, mice recovered from the cuprizone on a normal pellet diet.

### 2.5 Repeated in vivo imaging

Mice were imaged at baseline prior to starting the cuprizone diet, and then once every week while on the cuprizone diet and during recovery, out to 13 weeks. In a few mice, baseline imaging was performed immediately after chronic cranial window implantation, with the mouse remaining anesthetized during the transition from surgery to imaging. In most mice, imaging and cuprizone administration commenced about three weeks after window implantation. For imaging after window implantation, the mice were anesthetized with isoflurane (1.5-2% in oxygen) and given glycopyrrolate (0.5 mg/100 g; intramuscular) and 5% weight/volume (w/v) glucose in saline (100 μL per hour; subcutaneous). A few mice were injected with fluorescein (FITC)-conjugated dextran (5% w/v; 100 μL; retro-orbital; D1823, ThermoFisher) to help visualize the vasculature. Once anesthetized, mice were head-fixed in a stereotaxic apparatus and temperature controlled at 37°C with the feedback-controlled heating pad. Breathing rate was also recorded using a lab-built piezoelectric breathing sensor (Rivera et al., 2024) and isoflurane levels adjusted to ensure that breathing was stable at ∼60 breaths per minute.

Prior to the first multi-photon imaging session, a green-LED brightfield map of the cortical vasculature was acquired using a 4x objective (0.28 NA, Olympus). This map was used to record the surface starting location for navigating to the same subcortical WM areas during subsequent imaging sessions.

To acquire *in vivo* WM images, 3PEF and THG signals were excited and detected using the THG microscopy setup described above. After navigating to a location in the parietal cortex about 2 mm lateral from the sagittal sinus and free of larger surface vessels that can absorb some of the THG signal, the stage was raised to bring the laser focus deeper into the tissue until the WM became visible as bright bundles in the THG channel. This process was repeated until a suitable area was identified, and then stacks were acquired. Low magnification stacks were acquired at 2-µm steps through the WM and higher magnification stacks were acquired at 1-µm steps. At each z-position in the stack, three to ten frames were acquired and subsequently averaged to create one frame with higher image quality. Laser power was kept below ∼200 mW at the surface to avoid thermal damage.

### 2.6 Quantification of white matter degradation

Myelin integrity was evaluated in three different ways: blistering morphology, intranodal distance, and homogeneity of THG intensity. Based on literature and author observation, myelin blistering features were divided into four categories (blebs, swellings, vesicles, and holes) and their incidence counted in the image stacks. Intranodal distance was measured by manually identifying nodes and then using custom MATLAB script to quantify their length as the distance between inflection points of a four-term Gaussian fit of the average THG intensity profiles across each node (MATLAB fit() function with ‘gauss4’).

As a novel measure of myelin changes, a metric for evaluating the degree of blotching or mottling was developed using FIJI (Schindelin et al., 2012) and MATLAB. First, frames containing similar (or identical) representative myelin bundles were identified at each time point in FIJI and then a one-pixel median filter was applied to each image to remove impulse noise. The filtered images were then transferred to MATLAB, where a mask was made by thresholding each image. We then measured the size of the contiguous areas of pixels (or clusters) above the threshold using the MATLAB regionprops() function and plotted the cumulative probability distribution of the clusters. Shifts in the size of areas with above-threshold pixels lead to shifts in this distribution function, with more distributed and smaller areas typically observed in healthy control images and more clumping of above-threshold pixels into fewer, larger areas in cuprizone-treated mice. We empirically determined that a threshold value of the top 1% of pixels led to reliable differences in the cumulative probability distribution between control and cuprizone treated mice. The difference between cumulative distributions over time was represented as the difference between the integral of each distribution from cluster sizes of 1 to 50 pixels, using the MATLAB trapz() function. Code for this analysis and the intranodal distance measurement is available here: https://github.com/sn-lab/THG-myelin-manuscript.

### 2.7 Quantification of microglia involvement

The relationship between blistering events and microglia was quantified by calculating the Euclidean distance between them. The microglia bodies were extracted from the raw data by first applying a Gaussian blur to the GFP channel. The high intensity regions above an appropriate threshold, depending on the image quality, were then masked. The 3D Cartesian coordinate centroids of the resulting segmented microglia were measured and stored with FIJI. Centroids of blister-like events were also extracted with FIJI from manual ROI outlines and stored. For each identified blister-like event, the Euclidean distance to all microglia centroids were calculated and the distance to the nearest microglia was recorded with a custom MATLAB script. Edge effects were addressed by discarding nearest distances less than the boundary distance of the blister-like event. Then, for each voxel in the imaging volume, we calculated the Euclidean distance to the closest microglia centroid, again acknowledging edge effects. Distribution functions of these distances were generated to visualize the spatial distribution of microglia relative to blebs and to general voxels. Data from specific time points were analyzed to investigate temporal changes in the proximity of microglia to blister-like events. Code for this analysis is also available here: https://github.com/sn-lab/THG-myelin-manuscript.

### 2.8 Immunostaining for nodes of Ranvier

To validate nodes of Ranvier visible with THG signal, post-mortem tissue was labeled with traditional Caspr/Nav1.6 immunostaining (Poliak et al., 1999; Caldwell et al,. 2000). However, due to the necessary use of detergents to expose the proteins of interest for immunostaining, it was difficult to acquire high quality THG images of myelin from tissue that had already been stained, since the lipids in myelin are permeabilized by the detergent. Conventional methods of tissue preservation such as storing samples at 4°C in 10% sucrose or at -20°C in cryoprotectant solution also compromise the myelin structure and needed to be avoided. Therefore, all THG imaging was done as quickly as possible after perfusion and sectioning, as described below, and immunostaining protocols were performed afterwards and the images spatially aligned.

First, 5 minutes prior to perfusion, mice were retro-orbitally injected with 100 μL of Texas-Red-conjugated lectin (L32482, ThermoFisher) to label the vasculature. Then mice were perfused with 15 mL of cold 4℃ Krebs buffer (J67795.AP, ThermoScientific) and 40 mL of warm 37℃ 4% PFA (Blanke & Gray et al., 2024). Immediately after perfusion, the brains were removed and fixed in 4% PFA overnight at 4℃. The following morning, 4-5 holes were punched using 23-gauge blunt needle tips in each hemisphere along the superior-inferior axis to serve as fiducials for later image registration. Transverse sections of 30-µm thickness were sliced parallel to the cortical surface using a vibrating blade microtome (VT1200, Leica). Slices containing corpus callosum were immediately mounted on a glass slide, allowed to briefly dry, coverslipped with phosphate-buffered saline (PBS) as the mounting medium, and then imaged with THG/3PEF, averaging 5 frames.

After imaging, the coverslip was gently removed and the tissue was blocked in goat serum with Triton-X (9036-19-5, Sigma Aldrich). After 1 hour, the blocking buffer was removed and replaced with a primary antibody solution of 10% goat serum containing antibodies against Caspr (dilution 1:300; MABN69, Sigma-Aldrich) and Nav1.6 (dilution 1:200; #ASC-009, Alomone) and left to incubate at 4°C for 24 hours. At the completion of the primary antibody incubation, the tissue was again washed three times in PBS for 10 minutes each and then incubated for 1 hour with secondary antibodies: anti-mouse FITC for Caspr (dilution 1:200; 31569, Invitrogen) and anti-rabbit Texas Red for Nav1.6 (dilution 1:200; T2767, Invitrogen). Following secondary antibody incubation, the tissue was washed three more times in PBS for 10 minutes each and then coverslipped with Mowiol (#81381, Sigma-Aldrich) as the mounting medium for imaging on a confocal microscope (LSM710, Zeiss). The fiducial hole-punches were used to identify the same WM regions for image acquisition. Image stacks from THG and confocal were then manually registered using the fiducial hole-punches, matching autofluorescence, and 3PEF and linearly excited fluorescence signal from the lectin Texas-Red label (L32482, ThermoFisher). Confocal imaging data was acquired through the Cornell Institute of Biotechnology’s BRC Imaging Facility (RRID:SCR_021741).

### 2.9 Statistics

We made two statistical comparisons in this manuscript. For comparing nodal widths in Fig. 4, we used an unpaired t-test. For comparing the cumulative distribution functions in Fig. 5, we used the two-sample Kolmogorov-Smirnov test.

## 3 Results

### 3.1 High-resolution in vivo imaging of subcortical WM

Imaging through a chronic cranial window with 1320-nm femtosecond laser pulses consistently allows for parallel visualization of 3PEF from FITC in blood vessels and THG signals through the mouse cortex and into subcortical WM (Fig. 1A - C). Single, unaveraged image frames are sufficient to show that the texture of the THG signal changes dramatically with depth (Fig. 1D), from only single axons visible occasionally in the cortex to layers of compact, oriented bundles in the WM. At the bottom of the volume in Figure 1C, due to the presence of many aligned and thickly myelinated axons, the THG signal from the WM is readily identifiable as compared to that in the cortex where myelination is sparse and thinner (Basu et al., 2022). We also observed the expected change in the orientation of the axon bundles with depth through the WM. To improve image quality without causing thermal damage, frames were averaged instead of increasing laser power, enabling clear visualization of myelinated axons in the cortex (Fig. 1E) and bundles in the subcortical WM (Fig. 1F). Projecting across multiple frames in the z-direction (Fig. 1G) provided a more 3D representation of the bundles in the top layer of the corpus callosum (CC). The ability to resolve individual myelinated axons decreased with depth into the highly scattering CC (Fig. 1H). THG signal was also reduced by the presence of large surface blood vessels, which absorb much of the backscattered THG signal. Nevertheless, THG imaging at 1320 nm can routinely resolve micrometer-scale features in subcortical WM, opening the door to exploring dynamic changes in myelin in a disease model.

**Fig. 1.**
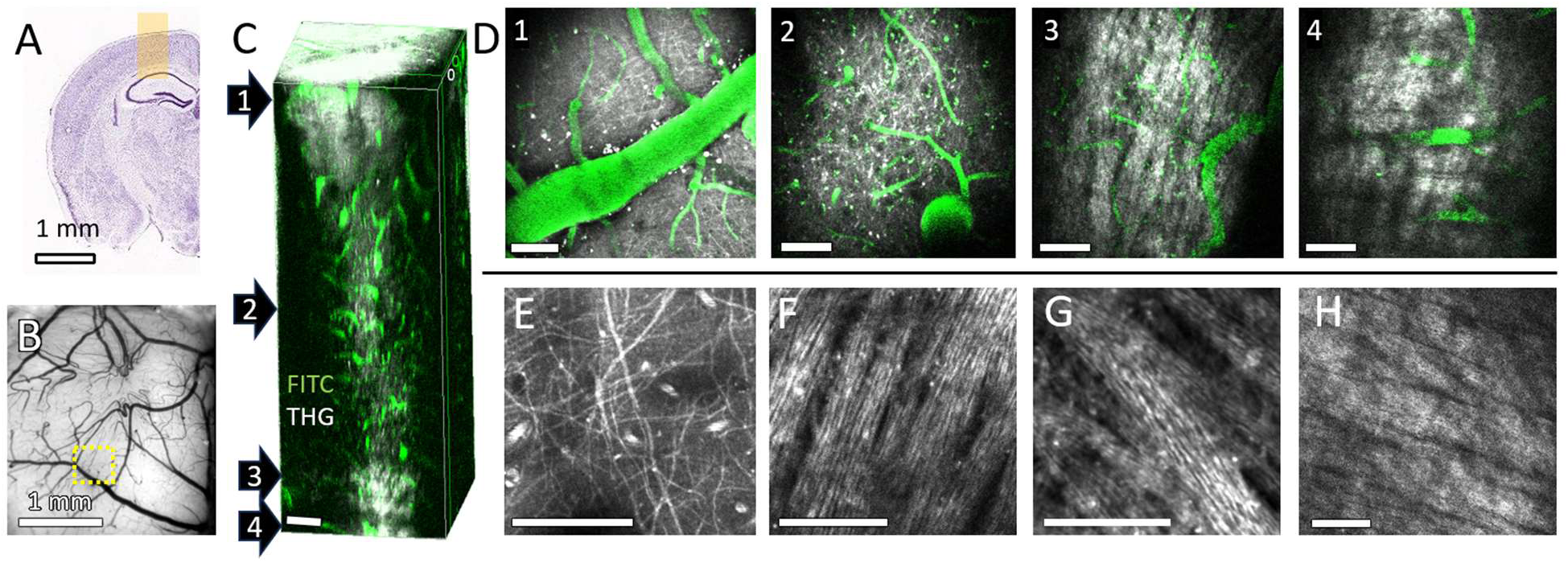
THG imaging at 1320-nm enables visualization of subcortical WM in mice. (A) Approximate location of imaging area, highlighted on position 281 from the Allen Brain Atlas (mouse.brain-map.org). (B) Green-light reflectance image of the cortex of a mouse under a cranial window. The location of the image stack in panel C is denoted with a yellow dotted outline. (C) 300x300x850 μm volume of THG and FITC-dextran (injected retro-orbitally) signal acquired from the top of mouse cortex down to the CC. The numbers to the left indicate the depths of the image frames shown in panel D. (D) Slices from the image stack in C at 50, 500, 800, and 850 μm deep into the mouse brain. (E) Frame-averaged THG from thin myelinated axons and red blood cells in thicker capillaries, 50 μm from the surface of the mouse cortex (20 frames over 2 μm). (F) Frame-averaged THG of bundles of myelinated axons at the surface of the CC, 710 μm deep (10 frames). (G) Z-projection through 5 µm thickness of a 1-µm spaced, frame-averaged (7 frames) THG image stack of myelinated axon bundles 770 μm deep. (H) Frame-averaged THG of large bundles of myelinated axons 40 μm below the surface of the CC, 790 μm deep from the surface of the cortex (5 frames). Scale bars = 50 μm, unless otherwise noted.

### 3.2 Longitudinal degeneration of myelin in cuprizone model

With a reliable method for visualizing myelin and the ability to image the same region by using surface vessels as a roadmap (Fig. 1B), it is possible to track myelin degeneration in a mouse model of disease over time. In the cuprizone model of MS, we imaged the same anatomically identified WM region over consecutive weeks of cuprizone administration and by the third or fourth week, changes in myelin structure were visible as blister-like events (Fig. 2A, B, Supplementary Vid. 1), which were not prevalent in controls (Supplementary Vid. 2). These blister-like events were previously described in humans with MS by Luchicchi et al (2021, 2024), while others have identified them in MS mouse models (Romanelli et al., 2016; Joost et al., 2022). Further, the THG image quality was sufficient to create classifications for several different types of myelin changes (Fig. 2C). Blisters within a myelin bundle were categorized as ‘blebs,’ which were morphologically distinct from the blisters between myelin bundles that we labeled as ‘vesicles,’ consistent with other nomenclature (Peters & Folger, bu.edu/agingbrain). We noted additional distinct features, including gaps that appear as myelinated axons splitting within a bundle that we termed ‘swellings,’ and openings that appear within or between bundles but without distinct myelin around the edge that we labeled ‘holes.’ With this THG imaging technique, all four types of morphological changes in myelin could be counted (Fig. 2E) and their size measured (Fig. 2F) across multiple weeks of cuprizone treatment (0 to 8 weeks) and recovery after cuprizone withdrawal (8 to 13 weeks). We found that by the third week of cuprizone treatment, there was already a marked increase in blister-like events, mostly blebs, which continued to increase over time during cuprizone administration (Fig. 2D, E). The number of pathological features began to decrease within a few weeks of cuprizone withdrawal (Fig. 2D). Interestingly, the distribution of the size of these blisters did not appear to change as more pathological features appeared during cuprizone administration and then disappeared after cuprizone withdrawal (Fig. 2F). We also quantified the density of putative lipid deposits (bright punctae present only in the THG channel, Fig. 2A, B), but did not detect a change after cuprizone administration.

**Fig. 2.**
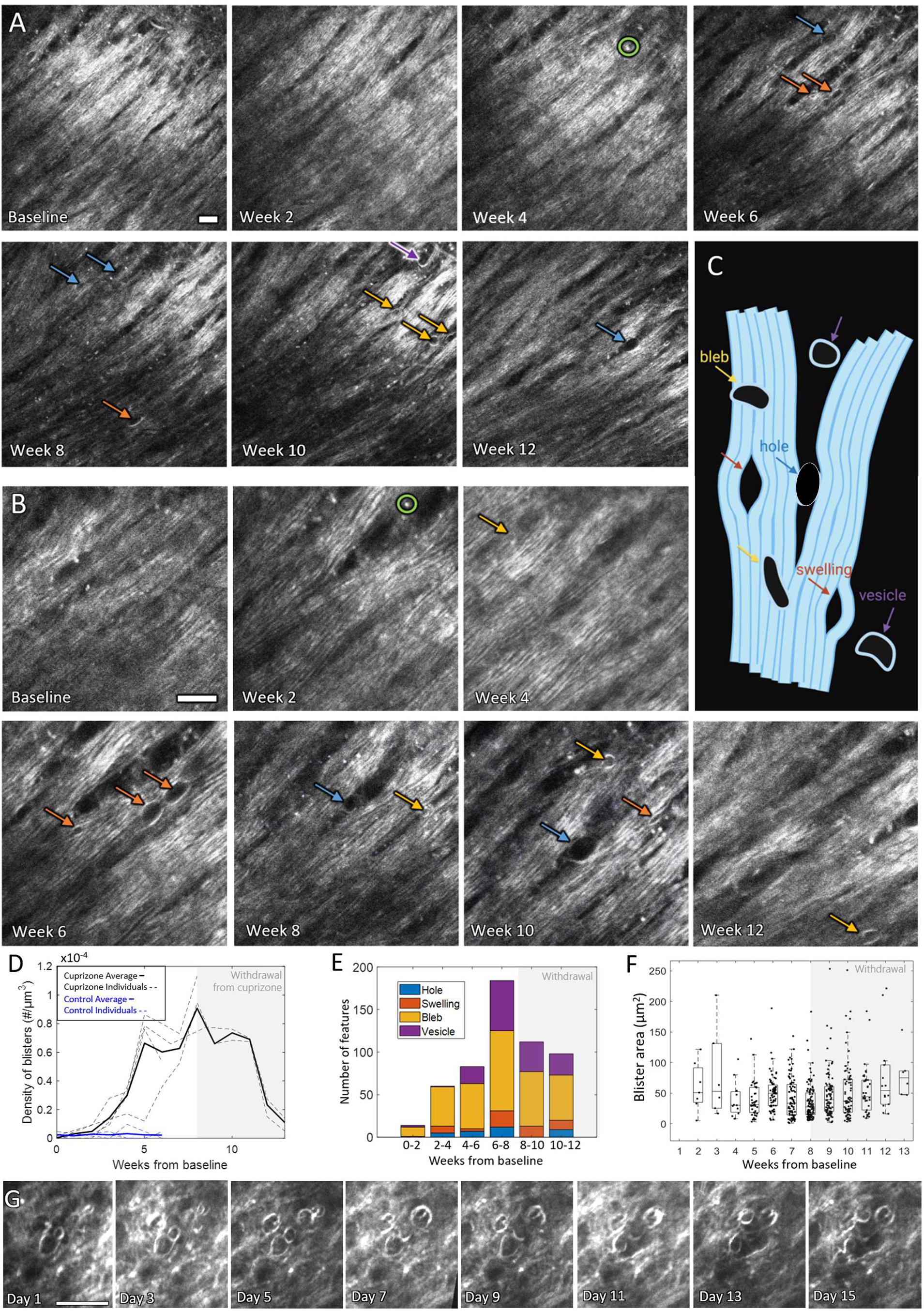
Visualization of subcortical myelin degeneration in mice receiving cuprizone. (A, B) Development of blistering in the myelin-axon unit with cuprizone administration (CX_3_CR-1^GFP^ mouse) at low and high level of magnification, respectively (3-5 frames averaged). Arrows indicate pathological blister-like events, with color-coding by type with guidance from literature (blue for holes, orange for swellings, yellow for blebs, and purple for vesicles). Green circles indicate examples of lipid deposits or lipofuscins. (C) Schematic of different categories of pathological blister-like events that were scored (created in BioRender. Hong, N. (2025) https://BioRender.com/z46dxie). (D) Plot of the density of blister-like features over time during cuprizone administration and withdrawal (shaded region). Trajectories for individual mice shown with dotted lines, while solid lines show the average (n = 4 and 3 for cuprizone and control, respectively). (E) Distribution of different types of pathological features over time. (F) Cross-sectional area of all blister-like events observed during cuprizone administration and withdrawal. (G) Tracking blister changes on alternate days revealing dynamic changes in myelin blisters after withdrawal from cuprizone (day 1 = 24 hours without cuprizone after 6 weeks of cuprizone administration; 10 frames averaged). Scale bars = 20 μm.

In addition to tracking overall changes in myelin morphology, THG imaging in the same location enabled tracking the dynamics of the same myelinated axon over time (Fig. 2G, Supplementary Fig. 1). Imaging daily or on alternate days is possible, granting access to subtle changes shown in Fig. 2G, where vesicles were observed to merge or deteriorate over the first two weeks of cuprizone withdrawal. While our focus was on characterizing changes in the subcortical WM, we observed similar blistering and degradation events in myelinated axons in the cortex (Supplementary Fig. 2).

Furthermore, THG imaging can be used in parallel with 3PEF to explore the interaction of different cell types with myelin changes. For example, imaging a Thy1-YFP mouse allows for *in vivo* study of the axon-myelin unit (Supplementary Fig. 3). Likewise, by imaging CX_3_CR-1^GFP^ mice, it was possible to quantify the degree of spatial association of microglia with myelin blisters during cuprizone administration (Fig. 3A). First, we compared the distribution of distances from each voxel in the image volume to the centroid of the nearest microglia cell body, taking care to avoid confounds from edge effects (Fig. 3B). We found the distribution of distances decreased with cuprizone administration (Fig. 3C), reflecting an expected increase in microglia density with cuprizone administration (Gudi et al., 2014), which was also evidenced by a ∼50% increase in area coverage of microglia in the CC. Next, we used the same technique to compare the voxel-to-microglia distance distributions to the bleb-to-microglia distance distributions. We found no difference between the two distributions after cuprizone administration or withdrawal, suggesting that microglia cell bodies are not associated with blebbing events.

**Fig. 3.**
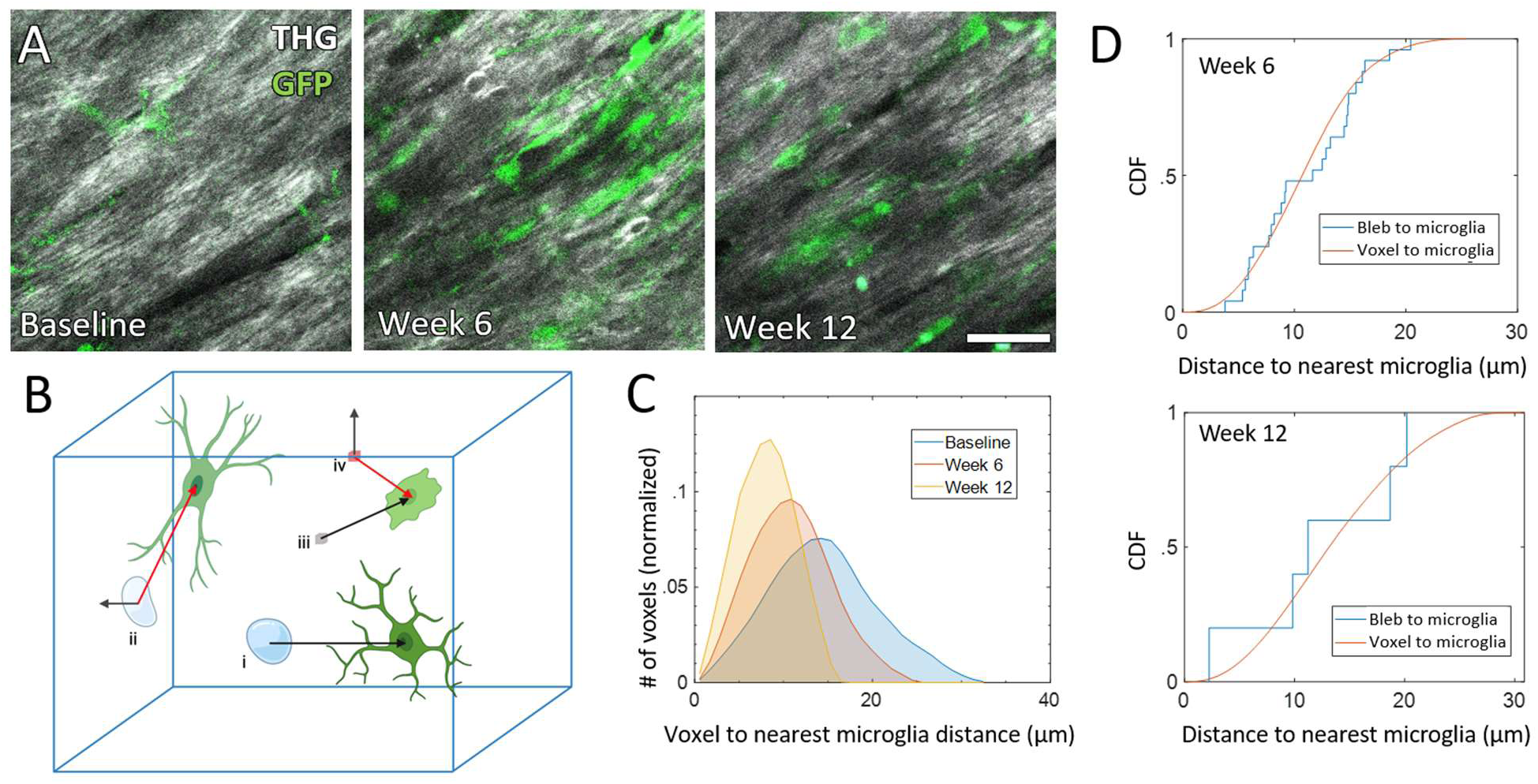
Microglia cell bodies do not associate with forming myelin blebs in the WM after cuprizone administration. (A) Visualizing microglia involvement in response to cuprizone in CX_3_CR-1^GFP^ mice (microglia shown in green; THG in greyscale; 3-4 frames averaged; scale bar = 20 μm). (B) 3D schematic of cases to determine inclusion or exclusion of voxels/blebs from analysis, wherein bleb (i) and voxel (iii) are included because they are closer to a microglia centroid than the edge of the volume, whereas bleb (ii) and voxel (iv) are excluded because they are closer to the edge of the volume than a microglia centroid (created in BioRender. Hong, N. (2025) https://BioRender.com/z46dxie). (C) Distribution of distances of each voxel to nearest microglia before cuprizone treatment (baseline), after six weeks of cuprizone treatment (week 6), and at eight weeks of cuprizone treatment followed by four weeks of withdrawal (week 12). (D) Cumulative distribution of microglia proximity to blister-like blebbing events, as compared to random locations, at two different timepoints.

### 3.3 In vivo measurement of width of nodes of Ranvier

A well-known first-stage marker of demyelination in MS and other diseases is an increase in the length of nodes of Ranvier, referred to as the intranodal distance (Arancibia-Carcamo & Attwell, 2014; Gallego-Delgado et al., 2020; Luchicchi et al., 2021). The image quality of the myelin bundles acquired with frame-averaging was sufficient to detect small decreases in signal intensity, on the order of one to two micrometers, along the length of axons in the THG image (Fig. 4E). We compared the length of putative nodes of Ranvier from THG imaging (Fig. 4A) with the intranodal distance measured using traditional Caspr-Nav1.6 antibody staining (Fig. 4B) for the same nodes of Ranvier in extracted, fixed subcortical tissue (Fig. 4C). We found a strong correlation between the measured intranodal distance using THG and antibody staining (Fig. 4C). Next, we measured the intranodal distance in putative nodes of Ranvier from *in vivo* THG imaging in healthy mice (*n* = 9) and in mice after 6-8 weeks of cuprizone administration (*n* = 3) (Fig. 4E and F). On average, we found that cuprizone-treated mice had a significantly larger intranodal distance (Fig. 4G; t-test, p < 0.0001).

**Fig. 4.**
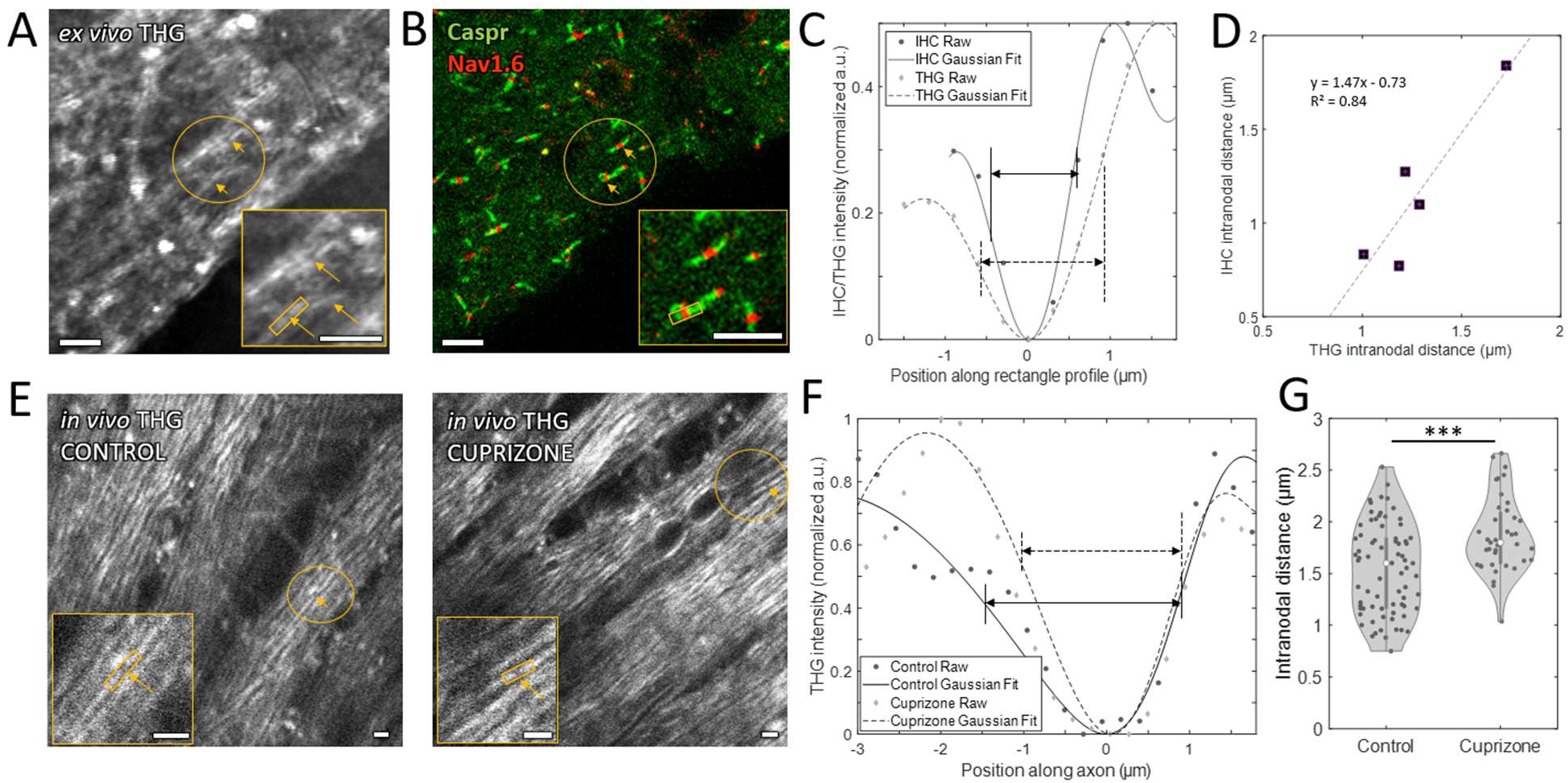
THG microscopy enables measurement of the intranodal distance in putative nodes of Ranvier, *in vivo*. Validation of node of Ranvier identification and measurement with immunostaining by identifying the same nodes with (A) *ex vivo* THG microscopy (5 frames averaged) and (B) confocal microscopy. (C) Line profiles of THG and Caspr staining of the node indicated by yellow rectangles in A and B. (D) Correlation of measured intranodal distance using THG and confocal imaging of Caspr staining (R^2^ = 0.84). (E) Identification of putative nodes of Ranvier *in vivo* in subcortical WM in control and cuprizone-treated mice (3-4 frames averaged). (F) Average of line intensity profiles across nodes indicated by yellow rectangles in E. (G) Measurement of intranodal distance in control and cuprizone-treated mice (***: p < 0.0001, t-test). Scale bars = 5 μm.

### 3.4 Early stages of demyelination visible with THG hyperintensities

While the ability to track the dynamics of myelin blister-like events in the subcortical WM is novel and important for understanding disease progression, there is significant interest in identifying earlier and more subtle markers of myelin changes, like the node of Ranvier length measurement described above. In the cuprizone model, we were able to use THG imaging to observe mottling or a blotchy texture in the myelin bundles as soon as two to three weeks on the cuprizone diet (Fig. 5A). We quantified this change in texture by analyzing the spatial distribution of the brightest THG signal. In line with our qualitative assessment, we found that the brightest pixels in the THG image became more spatially concentrated after two to three weeks of cuprizone. We quantified this by calculating the cumulative probability distribution of the number of pixels in clusters created from thresholding the 1% brightest pixels and comparing this over time (Fig. 5B). We found a significant change, relative to baseline, for mice on cuprizone (KS test; p < 0.005; n = 3 in each group; 2 to 3 weeks of cuprizone treatment), which was not observed in the control mice (Fig. 5C).

**Fig. 5.**
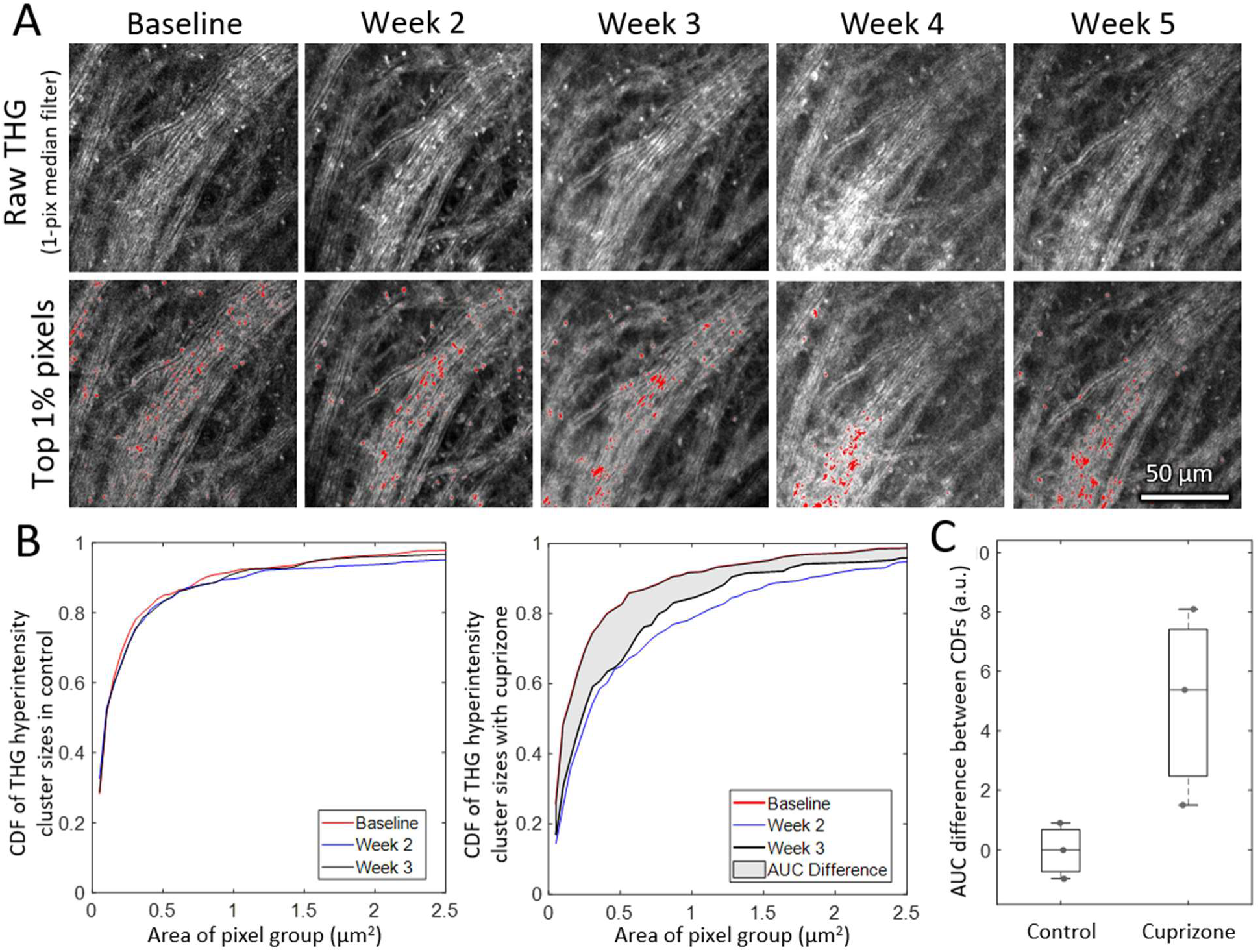
THG imaging of early myelin changes after cuprizone administration, measured by the congregation of the brightest signal pixels. (A) Mottling of the texture of THG images of WM bundles detected early in cuprizone administration. Top shows the raw signal, while the bottom shows the brightest 1% of pixels in red (4-6 frames averaged; scale bar = 50 μm). (B) Cumulative distribution function (CDF) of the size of contiguous pixel clusters that contain the brightest 1% of pixels in the image for control (left) and cuprizone treated (right) mice at baseline and at weeks 2 and 3 (area-under-curve (AUC) difference: control, none; cuprizone, KS test, p < 0.005 for week 2 and 3 vs baseline). (C) Quantification of AUC difference, relative to baseline, between CDFs for control and cuprizone-treated mice after 2-3 weeks.

## 4 Discussion

By using THG microscopy with 1320-nm excitation, we have demonstrated the ability to visualize myelinated axons at high resolution in the subcortical WM of live mice through the intact cortex. In a disease state induced by cuprizone administration, we were able to detect degenerative changes in the myelin structure. After imaging the same region across multiple weeks, we could track the emergence and increased incidence of myelin blistering events, as well as categorize these pathological events based on appearance. We further measured putative nodes of Ranvier as the width of small decreases in the myelin THG signal, a method that we confirmed by post-mortem correlation between the THG signal and fluorescent antibody staining. Finally, we observed subtle early changes in the myelin structure – before the appearance of the more obvious blistering events – that we quantified as THG hyperintensities

Compared to other approaches for *in vivo* imaging of WM, our THG microscopy approach uniquely enables label-free live imaging of myelin with cellular-level resolution at depths reaching to the subcortical WM in mice. While MRI and CT can visualize the WM through the whole brain – in mice and in humans – neither technique has the microscopic resolution needed to uncover underlying cellular mechanisms of myelin-related disease. CARS, on the other hand, is a popular technique for visualizing myelin degeneration. By using two laser beams to excite molecular vibrations, CARS is particularly sensitive to lipid-rich structures, like the myelin sheath (Yu et al., 2014). A related technique, SRS, removes the nonresonant background that impacts CARS, enabling high quality *in vivo* imaging of myelin with submicrometer resolution and facilitating measurement of quantitative metrics including g-ratio (Wang et al., 2005; Yu et al., 2014). However, CARS and SRS setups are much more complicated than standard multiphoton microscopes, and typically use shorter wavelengths for laser excitation, as compared to the 1320 nm used here, limiting depth penetration. Similarly, SCoRe provides quantitative information about myelin thickness and integrity (Schain et al., 2014; Craig et al., 2024), but depth penetration for this linear optical measurement is again limited to the mouse cortex. Meanwhile, 3PEF microscopy at 1320-nm excitation in transgenic mice with fluorescently-labeled myelin could enable some structural analysis, but this technique is still suboptimal since proteins in the myelin (e.g. myelin basic protein) are fluorescently tagged rather than the myelin itself.

THG microscopy, however, overcomes these limitations for *in vivo* imaging of myelin. Previous work in our group and others has capitalized on intrinsic THG signal from myelin to image WM in the mouse spinal cord (Farrar et al., 2011; Lim et al., 2014) and myelinated axons in the cortex (Redlich & Lim, 2019). While effective at obtaining high-resolution images of myelin in the spinal cord WM, 1030-nm excitation did not have sufficient penetration depth to reach the subcortical WM in the brain (Farrar et al., 2011). Similarly, Redlich and Lim’s work (2019) with 1160-nm excitation allowed quantification of myelination in the mouse cortex, including g-ratio measurements, but could not be applied at greater depths. Another study used polarized THG microscopy to quantify details of myelin organization around axons, but also using 1100-1150-nm excitation that limited depth penetration (Morizet et al., 2025). Recent work, however, has demonstrated the ability to use 1320-nm excitation to successfully reach the subcortical WM (Horton et al., 2013; Ouzounov et al., 2019) while avoiding the pathological artifacts caused by removing the cortex to optically access the WM. Here, we used this precedent and optimized the technique to study morphological changes of myelin in the subcortical WM. We note that THG is not molecularly specific to myelin, so other structures like blood vessels and lipid deposits are visible (Ahn et al., 2020; Small et al., 2017); however, these are structurally distinct.

The cuprizone mouse model of MS was an ideal imaging test because the demyelination process is well-studied with an overt pathology. In humans, one primary characteristic of myelin degeneration in MS is the formation of blisters or blebs (Lassmann, 2018; Luchicchi et al., 2021). In mice, Romanelli et al. (2016) reports ‘myelinosome’ formation during *in vivo* imaging of oligodendrocytes in the spinal cord of the experimental autoimmune encephalomyelitis mouse model of MS, and Joost et al. (2022) describes ‘vacuole’ formation, characterized histologically in the cuprizone mouse model of MS. However, these studies were unable to capture longitudinal demyelination dynamics in the subcortical WM, a primary location of pathological changes in MS and neurodegenerative diseases (Festa et al., 2024). Our THG imaging technique successfully visualized these blisters and also allowed for tracking of blister dynamics in the subcortical WM that is unachievable with any other current imaging techniques. By navigating to the same myelin bundles at each imaging session during cuprizone administration and withdrawal, we could measure the appearance, disappearance, and changing morphology of individual blisters.

Previous work has shown that nodes of Ranvier increase in width in MS (Arancibia-Carcamo & Attwell, 2014; Gallego-Delgado et al., 2020), so our result of a modest increase in intranodal width after cuprizone administration is not surprising. However, it is important to note that there is currently no robust technique to measure nodes of Ranvier *in vivo*, particularly in the subcortical WM. Since increased intranodal distance is an early marker of WM changes in MS and other neurodegenerative diseases (Arancibia-Carcamo & Attwell, 2014), the ability to measure nodes of Ranvier in mouse models could facilitate understanding of the causes and consequences of these changes.

In addition to showing we could characterize known myelin changes after cuprizone with THG imaging, we also developed a new THG-based metric based on the qualitative impression of blotchiness and mottling in the WM images. In previous work, we showed that the intensity of the THG signal from a slab of lipid surrounded by water increases exponentially with slab thickness, up to a thickness of 1 µm (for the tight focusing assumed in the model) (Farrar et al., 2011). Myelin can undergo an early degenerative change called decompaction, in which the protein anchors of successive myelin wraps weaken, leading to less tight packing and slight increases in myelin thickness (Gutiérrez et al., 1995; Giacci et al., 2018; Oost et al., 2023). The spatial clumping of the brightest THG signal that we detected in the early weeks of cuprizone administration is consistent with spatially-localized pockets of this myelin decompaction process, wherein more decompacted myelin produces brighter THG, due to increases in thickness compared to adjacent compact myelin and the exponential dependence of THG signal strength on thickness.

Finally, we confirmed the ability to correlate multiphoton imaging of cell behavior with myelin changes. Imaging transgenic mice with fluorescently tagged microglia allowed us to observe the increase in density of microglia in the damaged WM, and by collecting THG and 3PEF signals we were able to quantify their spatial changes as they related to the formation of blistering events. While we found no correlation between the locations of microglia cell bodies and blebs, future work can be extended to include microglia processes to fully capture any microglia involvement in the myelin degeneration events that are visible with THG. Pairing 3PEF imaging of other fluorescently labeled cell types with THG imaging will also allow further investigation of the interactions between cells of interest and myelin pathology.

## Acknowledgements

We thank Michael Lamont for building and maintaining our three-photon microscope and Tim Vartanian for commenting on a draft of this manuscript. We are also grateful to Irving Bigio, Alexander Gray, and Rhiannon Robinson for advice on preserving myelin integrity for post-mortem imaging. We acknowledge the Biotechnology Resource Center Imaging Core (RRID:SCR_021741) at Cornell for their help with confocal microscopy. This work was supported by NIH grants NS104350, EB002019, and NS126467.

