## Supplemental Table and Figures for "Label-Free Tracking of Subcortical White Matter Degradation *In Vivo* Using Third Harmonic Generation Microscopy in a Mouse Model of Multiple Sclerosis"

**Supplementary Table 1.** Breakdown of mouse genotypes and sex used by figure.

| Figure panel | Mouse genotype(s) (#; sex) |
|---|---|
| Fig. 1B-D | C57BL/6 (1; F) |
| Fig. 1E-H | ApoE3-TR (1, 3; F, M) |
| Fig. 2A, B; Supp. Vid. 1 | $CX_3CR\text{-}1^{GFP}$ (1; F) |
| Supp. Vid. 2 | $CX_3CR\text{-}1^{GFP}$ (1; F) |
| Fig. 2D-F | Control: C57BL/6 (2; M), $CX_3CR\text{-}1^{GFP}$ (1; F); Cuprizone: PLP-eGFP (1; M), $CX_3CR\text{-}1^{GFP}$ (1; F), ApoE3-TR (1; F) |
| Fig. 2G | Thy1-YFP (1; F) |
| Fig. 3 | $CX_3CR\text{-}1^{GFP}$ (1; F) |
| Fig. 4E, F | $CX_3CR\text{-}1^{GFP}$ (2; F) |
| Fig. 4G | Control: ApoE3-TR (3, 6; F, M); Cuprizone: ApoE3-TR (2; F), $CX_3CR\text{-}1^{GFP}$ (1; F) |
| Fig. 5A | ApoE3-TR (1; F) |
| Fig. 5B, C | Control: C57BL/6 (2; M), $CX_3CR\text{-}1^{GFP}$ (1; F); Cuprizone: Thy1-YFP (1; F), $CX_3CR\text{-}1^{GFP}$ (1; F), ApoE3-TR (1; F) |
| Supp. Fig. 1 | ApoE3-TR (1; F) |
| Supp. Fig. 2 | ApoE3-TR (1; F) |
| Supp. Fig. 3 | Thy1-YFP (2; F) |

**Supplementary Video 1.** Compilation across weeks of cuprizone administration and withdrawal of z-stacks in the same anatomically identified WM region (3-5 frames averaged). Blister-like events are visible as early as week 2. Scale bar = 20 μm.

**Supplementary Video 2.** Control dataset shown as a compilation across weeks of THG imaging of z-stacks in the same anatomically identified WM region (3-6 frames averaged). Obvious changes in myelin morphology, like blistering, are not prevalent. Scale bar = 20 μm.

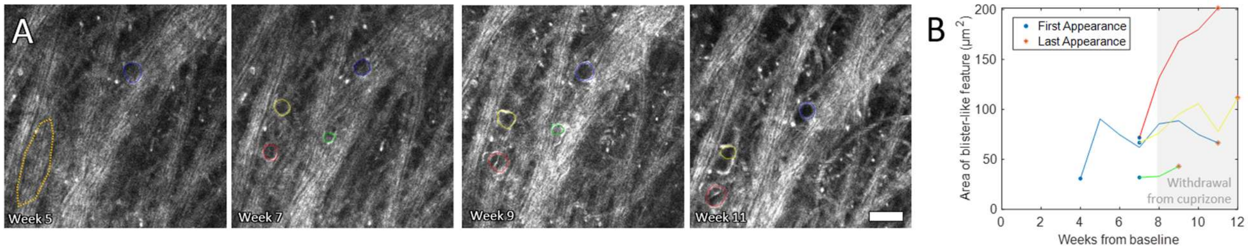


**Supplementary Fig. 1. Tracking individual blistering events during cuprizone administration and withdrawal.** (A) Pathological features in the CC are circled with colors corresponding to the traces of their size (B) after several weeks of cuprizone administration and withdrawal. Two of the four blister-like features tended to grow, even after cuprizone withdrawal. In the left-most panel, a degrading bundle is also outlined in dark yellow. Scale bar = 20 μm.

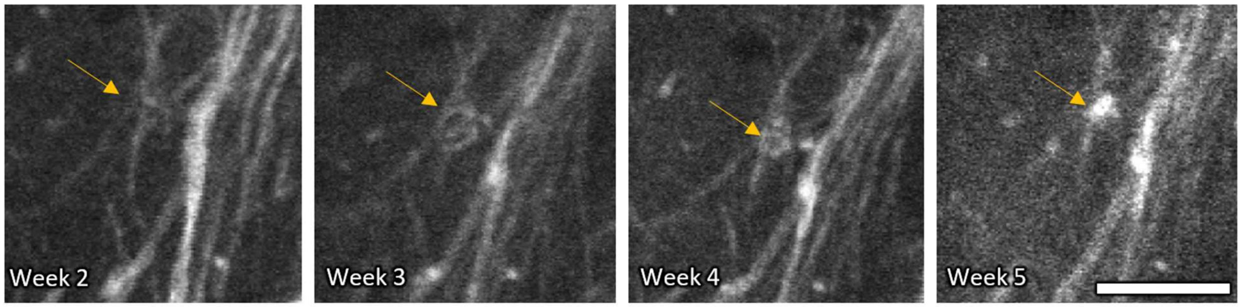


**Supplementary Fig. 2. Tracking individual blistering event in the mouse cortex during cuprizone administration.** Yellow arrow indicates a myelinated axon that starts blistering by three weeks of cuprizone administration and degrades into a bright lipid deposit by the fifth week. Scale bar = 20 μm.

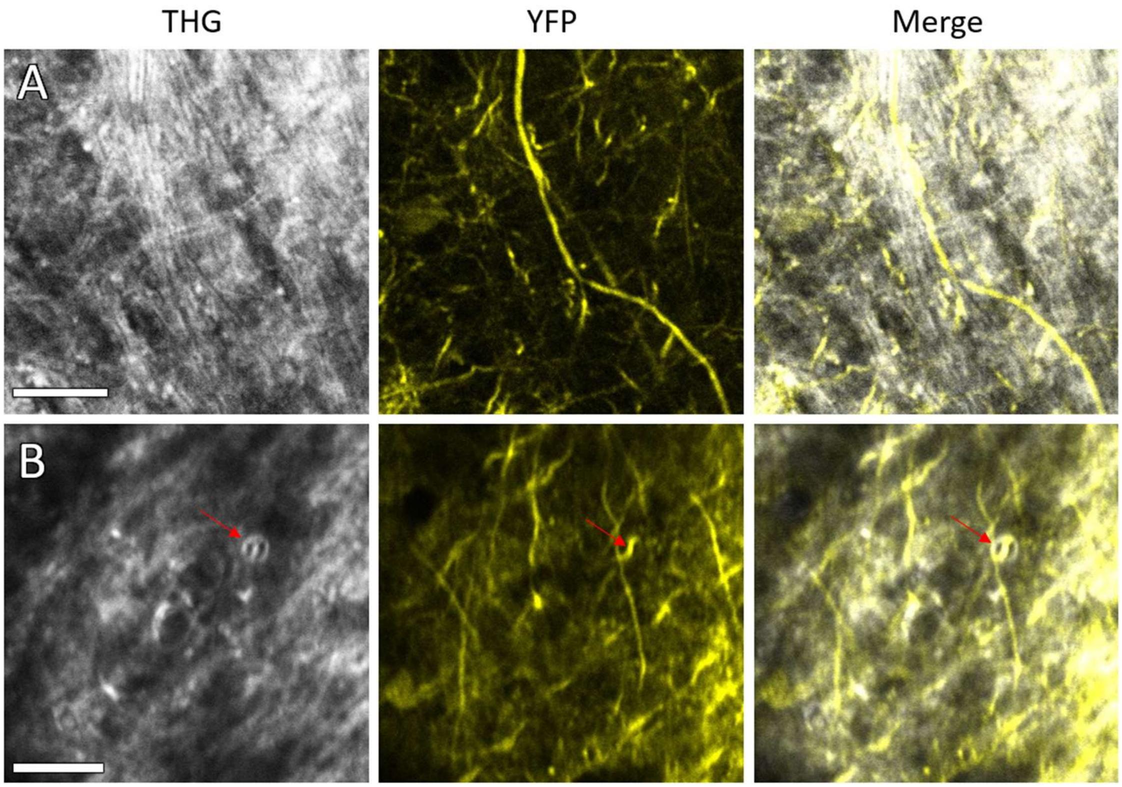


**Supplementary Fig. 3. Visualizing the axon-myelin unit in a Thy1-YFP mouse.** (A) Large axon in the CC (traversing center top to lower right) with myelin visible by THG surrounding the axon. (B) Pathological event in deep cortex after cuprizone administration. Red arrow indicates an axon within a myelin blister. Scale bars = 30 μm.
